# Effects of elevated CO_2_ on predator avoidance behaviour by reef fishes is not altered by experimental test water

**DOI:** 10.1101/050062

**Authors:** Philip L. Munday, Megan J. Welch, Bridie J.M. Allan, Sue-Ann Watson, Shannon McMahon, Mark I. McCormick

**Affiliations:** ARC Centre of Excellence for Coral Reef Studies, James Cook University, Townsville, QLD 4811, Australia; College of Marine and Environmental Science, James Cook University, Townsville, QLD 4811, Australia

**Keywords:** ocean acidification, carbon dioxide, antipredator behaviour, larval fish, GABA

## Abstract

Pioneering studies into the effects of elevated CO_2_ on the behaviour of reef fishes often tested high-CO_2_ reared fish using control water in the test arena. While subsequent studies using rearing treatment water (control or high CO_2_) in the test arena have confirmed the effects of high CO_2_ on a range of reef fish behaviours, a further investigation into the use of different test water in the experimental arena is warranted. Here, we used a fully factorial design to test the effect of rearing treatment water (control or high CO_2_) and experimental test water (control or high CO_2_) on antipredator responses of larval reef fishes. We tested antipredator behaviour in larval clownfish *Amphiprion percula* and ambon damselfish *Pomacentrus amboinensis*, two species that have been used in previous high CO_2_ experiments. Specifically we tested if: 1) using control or high CO_2_ water in a two channel flume influenced the response of larval clownfish to predator odour, and 2) using control or high CO_2_ water in the test arena influenced the escape response of larval damselfish to a startle stimulus. Finally, 3) because the effects of high CO_2_ on fish behaviour appear to be caused by altered function of the GABA-A neurotransmitter we tested if antipredator behaviours were restored in clownfish treated with a GABA antagonist (gabazine) in high CO_2_ water. Larval clownfish reared from hatching in control water (496 μatm) strongly avoided predator cue whereas larval clownfish reared from hatching in high CO_2_ (1022 μatm) were attracted to the predator cue, as has been reported in previous studies. There was no effect of testing fish using control or high CO_2_ water in the flume. Larval damselfish reared for 4 days in high CO_2_ (1051 μatm) exhibited a slower response to a startle stimulus, slower escape speed and a shorter escape distance compared with fish reared in control conditions (464 μatm). There was no effect of test water on escape responses. Treatment of high-CO_2_ reared clownfish with 4 mg l^−1^ gabazine in high CO_2_ seawater restored the normal response to predator odour, as has been previously reported with fish tested in control water. Our results show that using control water in the experimental trials did not influence the results of previous studies on antipredator behaviour of reef fishes and also supports the results of novel experiments conducted in natural reef habitat at ambient CO_2_ levels.

## Introduction

Rising concentrations of atmospheric carbon dioxide (CO_2_) have caused an increased uptake of CO_2_ by the ocean, leading to a decline in seawater pH and changes in the relative concentration of carbonate and bicarbonate ions, a process called ocean acidification (Sabine et al., 2004: Doney, 2010). The partial pressure of carbon dioxide (pCO_2_) at the ocean surface is increasing at the same rate at the atmosphere (Doney, 2010) and thus marine species will need to deal with rising CO_2_ levels in addition to declining pH and other changes in ocean chemistry (Pörtner et al., 2004). Recent models suggest an amplification of seasonal cycles in ocean pCO_2_ as atmospheric CO_2_ continues to rise, such that surface ocean pCO_2_ will reach 1000 μatm in summer for atmospheric CO_2_ concentrations that exceed 650 parts per million. Hypercapnic conditions (>1000 μatm CO_2_) are projected occur in up to half the surface ocean by 2100 on the current CO_2_ emissions trajectory (McNeil & Sasse, 2016).

Elevated CO_2_ levels can have a variety of effects on the physiology, life history and behaviour of marine organisms (Pörtner et al., 2004). Recent studies have demonstrated that high CO_2_ levels can fundamentally alter the behaviour of marine fishes and some invertebrates (reviewed by Briffa et al., 2012; Clements & Hunt, 2014; Heuer & Grosell, 2014; Nagelkerken & Munday, 2016). Of particular concern is an impaired response to the threat of predation, because it may increase mortality rates in natural populations (Munday et al., 2010; Ferrari et al., 2011a; Chivers et al., 2014a). Larval and juvenile reef fish exhibit an innate aversion to predator odour and chemical alarm cues from injured conspecifics because they indicate a heightened risk of predation (Holmes et al., 2010; Dixson et al., 2012). Individuals typically respond to these olfactory cues by reducing activity and seeking shelter (Ferrari et al., 2010). However, larval reef fish that have been reared at near-future CO_2_ levels do not reduce activity or stop feeding in the presence of chemical alarm cues (Ferrari et al., 2011a; Lönnstedt et al., 2013) and even become attracted to predator odour and chemical alarm cues when presented in a two channel flume (Dixson et al., 2010; Welch et al., 2014). Furthermore, juvenile fish exposed to high CO_2_ lose their ability for associative learning (Ferrari et al., 2012), a process that enables fine-tuning of risk assessment by the association of chemical alarms cues with the identity of specific predators. Finally, high CO_2_ affects the kinematics of predator-prey interactions, with juvenile prey exposed to elevated CO_2_ allowing predators to get closer before responding (Allan et al., 2013) and having reduced escape speeds and distances compared with fish reared at current-day CO_2_ levels (Allan et al., 2013, 2014). These changes in behaviour alter the outcome of predator-prey interactions, leading to significantly increased rates of mortality of small juveniles in mesocosm experiments (Ferrari et al., 2011b) and in fish transplanted to natural coral reef habitat (Munday et al., 2010; 2012; Chivers et al., 2014a).

Many of the pioneering studies on the effects of elevated CO_2_ on reef fish behaviour (e.g. Munday et al., 2009, Dixson et al., 2010; Ferrari et al., 2011a,b; Cripps et al., 2011) involved rearing fishes in control and elevated CO_2_ conditions and then testing their responses to olfactory cues in control water only. This method was chosen in these early experiments owing to logistical constraints and because pilot experiments showed that the response of high CO_2_ exposed fish to predator cue did not differ if the cue was presented in either control or high CO_2_ water (Dixson et al., 2010; Munday et al., 2010). Moreover, studies with freshwater fishes had shown that a pH reduction of 0.5 units in freshwater (<pH6.5) irreversibly alters the structure of chemical alarm cues and can dramatically affect prey antipredator responses (Brown et al., 2002; Leduc et al., 2004a,b; reviewed in Leduc et al., 2013). Consequently, there was concern that testing marine fish in CO_2_-acidified water (albeit still above pH7.5) could confound the interpretation of any effects of high CO_2_ on the behavioural response of the fish with a diminished efficacy of the olfactory cue itself. Indeed, subsequent studies have shown that chemical alarm cues from reef fishes degrade much more quickly in high CO_2_ water than they do in current-day conditions (Chivers et al., 2014b); therefore, experiments testing olfactory responses of marine species to high CO_2_ need to take this into account. Finally, small-scale laboratory experiments often have limited ecological relevance and it is challenging to extrapolate their findings to natural communities and ecologically relevant spatial scales (Naglekerken & Munday, 2016). Consequently, it was critical to test the effects of exposure to high CO_2_ in natural habitats in the field. This involved transplanting larval fish that had been reared in either control or high CO_2_ seawater back into natural coral reef habitat and monitoring their behaviour (Munday et al., 2010, 2012; McCormick et al., 2013).

Since these initial studies, a number of other experiments have been conducted in which the respective treatment seawater in which the fish were reared (control or high CO_2_) has been used in the experimental trials, confirming the results of earlier studies (e.g. Allan et al., 2014; Dixson et al., 2014; Welch et al., 2014; Nagelkerken et al., 2015) and/or revealing other behavioural effects of high CO_2_, such as an impaired response to auditory cues (Simpson et al., 2011). While previous pilot studies found no effect of the test water on the response of high-CO_2_ fish in behavioural assays (Dixson et al., 2010; Munday et al., 2010) and subsequent studies conducted in the respective treatment water have produced confirmatory results (Allan et al., 2014; Welch et al., 2014; Dixson et al., 2015), a more thorough validation study of the test water used in antipredator experiments is warranted. Here, we compared the use of control versus high CO_2_ seawater (called test water from hereafter) in experimental trials designed to test the antipredator responses of larval reef fishes reared in high CO_2_. We conducted antipredator experiments for two species that have been widely used in previous high CO_2_ experiments, the clownfish *Amphiprion percula* and the ambon damselfish *Pomacentrus amboinensis*. Specifically we tested if: 1) using control or high CO_2_ water in a two channel flume influenced the response of larval clownfish to predator odour in fish reared in either control or high CO_2_, and 2) using control or high CO_2_ water in the test arena influenced the kinematic response of larval damselfish to a startle stimulus in fish reared in either control or high CO_2_. Finally, because the effects of high CO_2_ on fish behaviour appear to be due to altered function of GABA-A neurotransmitter receptors (Nilsson et al., 2012; Hamilton et al., 2013; Heuer & Grosell, 2014) we tested if antipredator behaviours were restored in clownfish treated with a GABA antagonist (gabazine) in high CO_2_ water. Previous studies have demonstrated that abnormal antipredator behaviour of clownfish and damselfish reared in high CO_2_ conditions is reversed following treatment with gabazine (Nilsson et al., 2012; Chivers et al., 2014a). However, these previous studies involved high CO_2_ fish that were treated with gabazine in control water and then behaviourally tested in control water. In this experiment we therefore conducted the same gabazine treatments as previously used, except the drug was administered and behaviour was tested in high CO_2_ water.

## Methods

The experiment with larval clownfish was conducted at James Cook University’s (JCU) experimental aquarium facility in Townsville, Australia, where previous high CO_2_ experiments with this species have been conducted. The experiment with larval ambon damselfish was conducted at Lizard Island Research Station on the northern Great Barrier Reef, where previous high CO_2_ experiments with this species have been conducted. Both experiments were conducted during February 2016. Because our goal was to assess the use of control seawater in experimental trials of previous experiments (e.g. Dixson et al., 2010; Munday et al., 2010, 2012) we used the same methods as previous experiments, except that we applied a fully factorial design where fish reared in control and high CO_2_ conditions were behaviourally tested in both control and high CO_2_ water.

### Response to predator cue

Two clutches of larval clownfish from different parental pairs maintained at JCU were reared in control and high CO_2_ conditions from hatching until settlement (11-12 days post hatching) using standard practices (Wittenrich, 2007). Briefly, larvae were reared in 100 l incubator tanks supplied with a continuous flow of treatment seawater. Water temperature was maintained at 29 °C. Photoperiod was 12 hours light and 12 hours dark. A non-viable microalgae blend (Nano 3600 – Reed Mariculture™) was drip-fed into the incubator tanks during daylight hours to reduce light levels and enhance larval feeding during the first four days. Larvae were fed live rotifers (15 per ml) for the first four days and then transitioned onto artemia (5 per ml). Fish were not fed on the morning of testing. Each clutch was divided at hatching so that half the clutch was reared in control seawater (496 ± 55 S.D. μatm CO_2_) and half reared in the high CO_2_ treatment (1022 ± 37 S.D. μatm CO_2_). Two 10,000 L recirculating aquarium systems supplied seawater to the incubator tanks, one supplied control seawater and the other high CO_2_ seawater. To achieve the desired CO_2_ level in the high CO_2_ treatment, seawater was dosed with CO_2_ to a set pH in a 3000 L sump using an Aqua Medic AT Control System (Aqua Medic, Germany). The pCO_2_ in rearing tanks was measured by nondispersive infrared (NDIR) following the method described by Hari et al. (2008). Air in a closed loop was circulated by a small pump (11 min^−1^ flow rate) through a coil of thin-walled (membrane thickness ø 0.4 mm, outer ø 3.8 mm) medical grade silicone tubing placed in the tank. CO_2_ in the closed loop was then measured at 1 minute intervals with a Vaisala GMP343 infrared CO_2_ probe (accuracy ±5 ppm CO_2_ + 2% of reading over the range of experimental manipulations). Each reading (N=9 in control and N= 9 in high CO_2_) lasted 1 hour to ensure complete equilibration of CO_2_ between tank seawater and the closed loop of air. The average CO_2_ for the final ten minutes was calculated for each reading.

The response of settlement stage clownfish to predator cue was tested in a two-channel flume specifically designed for testing olfactory preferences of larval fish (Gerlach et al., 2007). Larvae were given the choice between a stream of seawater containing predator cue versus a stream of seawater without the predator cue. Both streams of water were either control or high CO_2_ test water for a given trial. Water from the respective paired sources was gravity fed into the flume at 100 ml min^−1^, maintained by a flow meter and dye test with every water exchange. Water was exchanged after every fish, with random alternation between control and high CO_2_ test water. For each trial, a single fish was placed in the centre of the downstream end of the flume and allotted a two-minute acclimation period. The position of the fish at 5-second intervals was recorded for the next two minutes. A one-minute rest period followed during which the water sources were switched to eliminate potential side preference. After the water switch, the fish was re-centred in the downstream end of the flume and the entire acclimation (2 minutes) and test period (2 minutes) were repeated. The temperature of the test water was kept within 1 °C of the water temperature in the rearing tanks. Three trials where the difference in water temperature between the flume and the rearing tanks exceeded 1 °C were discarded. The experimenter was blinded to the rearing treatment of the fish.

Predator cues were obtained from the common coral-cod, *Cephalopholis miniatus*. Five cod collected from the Great Barrier Reef (supplied by Cairns Marine, Cairns QLD) were maintained in 60 L aquariums supplied with a continuous flow of seawater from a different seawater system to that used to rear the clownfish. Each cod was fed one pellet of commercial fish food (Marine Food, Fish Fuel Co™ – 1 cube per fish) each evening. Water temperature was maintained at 29 °C. To obtain predator cues for the experiment, the water was drained from two predator tanks and refilled with water from the same system used to rear the clownfish. To match the methods used in previous experiments, water to the predator tank was turned off and predator cue water was collected after a 2 hours soak time. Predator tanks were aerated to maintain oxygen content and water-bathed to maintain a stable temperature. One of the two predator tanks was dosed with CO_2_ at the end of the 2 hour soak period, immediately before cue water was extracted. CO_2_ was dosed at low pressure by hand, with gentle stirring, until the pH matched that of the high CO_2_ rearing treatment. The other predator tank was also gently stirred but was not dosed with CO_2_. The individual cod used for predator cues was alternated within and between days. An identical procedure was followed with two tanks that did not house a predator to obtain seawater without predator cue to use in the flume.

#### Gabazine treatment

The gabazine treatment followed the flume methodology described above, except that fish were treated with 4 mg l^−1^ gabazine for 30 minutes immediately before testing in the flume. Eight larval clownfish were randomly selected from the high CO_2_ treatments and placed in 40 ml of high CO_2_ water containing gabazine. The fish were gently removed from the gabazine after 30 minutes and transferred to the flume using a small net. Their response to predator odour was tested in high CO_2_ water only. Previous experiments demonstrating that gabazine restores antipredator responses in larval fishes reared in high CO_2_ water have been conducted in control water (Nilsson et al., 2012; Chivers et al., 2014a). This experiment only tested if a similar effect was observed when gabazine was administered and fish tested in high CO_2_ water.

### Kinematics of escape response

Settlement stage larval *Pomacentrus amboinensis* were collected using light traps in the Lizard Island Lagoon and transferred to an environmentally-controlled aquarium facility at Lizard Island Research Station. Larval fish were assigned randomly to four replicate control (464 ± 32 S.D. μatm CO_2_) or four replicate high-CO_2_ (1051 ± 18 S.D. μatm CO_2_) aquaria. Twenty five *P. amboinensis* were housed in each 32 l (38Lx28Wx30H cm) aquarium. Water temperature was maintained at approximately 29.5 °C. Fish were kept for 4 days in treatment conditions and fed *Artemia* sp. twice daily *ad libitum*. Fish were not fed on the morning of testing. Each aquarium was supplied with control or elevated-CO_2_ seawater at 750 ml.min^−1^. Elevated-CO_2_ seawater was achieved by dosing with CO_2_ to a set pH, following standard techniques (Gattuso et al., 2010). Seawater was pumped from the ocean into 32 l header tanks where it was diffused with ambient air (control) or 100% CO_2_ to achieve the desired pH (elevated-CO_2_ treatment). A pH-controller (Aqua Medic, Germany) attached to the CO_2_ treatment header tank maintained pH at the desired level. Duplicate control and high CO_2_ systems supplied seawater to two replicate aquaria in each system. The pCO_2_ in rearing tanks was measured by NDIR as described above (N= 9 in control and N= 5 in high CO_2_).

After rearing for four days in control or high CO_2_ conditions, the escape response of the damselfish was tested in both control and high CO_2_ water (i.e. a fully crossed design). Individual fish were placed into the testing arena, which consisted of a transparent circular acrylic arena (diameter 200 mm) within a larger plastic tank (585 x 420 x 330 mm; 60 L) filled to a depth of 100 mm with either control or high CO_2_ water. A shallow water depth was used to limit vertical displacement of the fish during escape response trials. Water temperature in the experimental arena was 29°C - 30°C and the water treatment used was alternated after every 2^nd^ trial. The arena was illuminated by LED strip lighting (750 lumens) placed above the water surface on the outside of the tank. Five minutes after being released into the testing arena, an escape response was elicited by the release of a tapered metal weight from above the water surface. This was accomplished by turning off an electromagnet to which the metal weight was attached. The metal weight was controlled by a piece of fishing line that was long enough such that the tapered tip only just touched the surface of the water. In order to provide a sudden stimulation and allow quantification of escape latency, the stimulus was released through a white PVC tube (diameter 40 mm, length 550 mm) suspended above the experimental arena, with the bottom edge at a distance of 10 mm above the water level. Fish were only startled when they moved to the middle portion of the tank, allowing an individual to move an equal distance in any direction and standardising for fish position relative to the stimulus. Escape responses were recorded at 480 frames per second (Casio EX-ZR1000) as a silhouette from below obtained through pointing the camera at a mirror angled at 45°. To minimise visual disturbances, black sheeting surrounded the front of the mirror so that any movement within the room was undetected by the fish. Treatment water in the testing arena was changed every 2^nd^ trial to minimise CO_2_ loss in the high CO_2_ treatment water. The rate of CO_2_ loss from the testing arena was found to be negligible within this time frame. The experimenter was blinded to the rearing treatment of the fish.

From the videos, we quantified latency to initiate an escape, escape distance, escape speed and maximum speed. A 1 cm line was drawn in the centre of the inner arena to enable calibration for video analysis. All videos were analysed with the observer blind to the treatments.

#### Kinematic variables

Kinematic variables associated with the fast-start response were analysed using the image-analysis software Image-J, with a manual tracking plug-in. The centre of mass (CoM) of each fish was tracked for the duration of the response. The following kinematic variables were measured:

1. Latency to respond (sec) was measured as the time interval between the stimulus onset and the first detectable movement leading to the escape of the animal.
2. Escape distance (m) is a measure of the total distance covered by the fish during the first two flips of the tail (the first two axial bends, i.e. stages 1 and 2 defined based on Domenici and Blake (1997), which is the period considered crucial for avoiding ambush predator attacks (Webb, 1976).
3. Escape speed (m s^−1^) was measured as the distance covered within a fixed time (first 24 milliseconds after initial response) which corresponds to the average duration of the first two tail flips (the first two axial bends, i.e. stages 1 and 2 based on Domenici & Blake, 1997). This period is considered crucial for avoiding predator ambush attacks (Webb, 1976).
4. Maximum speed (m s^−1^) was measured as the maximum speed achieved at any time during stage 1 and stage 2.

### Data analysis

Two way ANOVA was used to compare percent time that larval clownfish spent in predator cue in the flume. Percentage data was logit-transformed for analysis (Warton & Hui, 2001). Treatment water (control or high CO_2_) and test water (control or high CO_2_) were the fixed factors. For the gabazine experiment, the percentage time that larval clownfish treated with gabazine spent in predator cue in the flume was compared with control fish using a t-test on logit-transformed data. A linear mixed effects model (LME) was used to compare escape responses of larval damselfish. Latency, escape distance, escape speed and maximum speed were each tested with treatment and test water as fixed factors. Tank was included in each model as a random effect to account for the subsampling of fish from replicate tanks within each CO_2_ treatment. The normality, linearity and homoscedasticity of residuals of the models were verified by visual inspection of residual-fit plots. Latency data was log10 transformed before analysis to improve the distribution of the data. ANOVA was conducted with Statistica (version x) and LMEs were fitted using the ‘nlme’ package in the R software package (Pinheiro et al. 2013).

## Results

There was a highly significant effect of treatment water on the percentage time that larval clownfish spent in the water stream containing predator odour (F_1,57_=6.256, p = 0.015), but no effect of test water (F_1,57_=0.0003, p = 0.973). Laval clownfish reared in control water strongly avoided predator odour, spending <9% of the time in that water stream (Figure 1). In contrast, larval clownfish reared in high CO_2_ water were strongly attracted to predator odour, spending nearly 80% of the time in that water stream (Figure 1).

**Figure 1.**
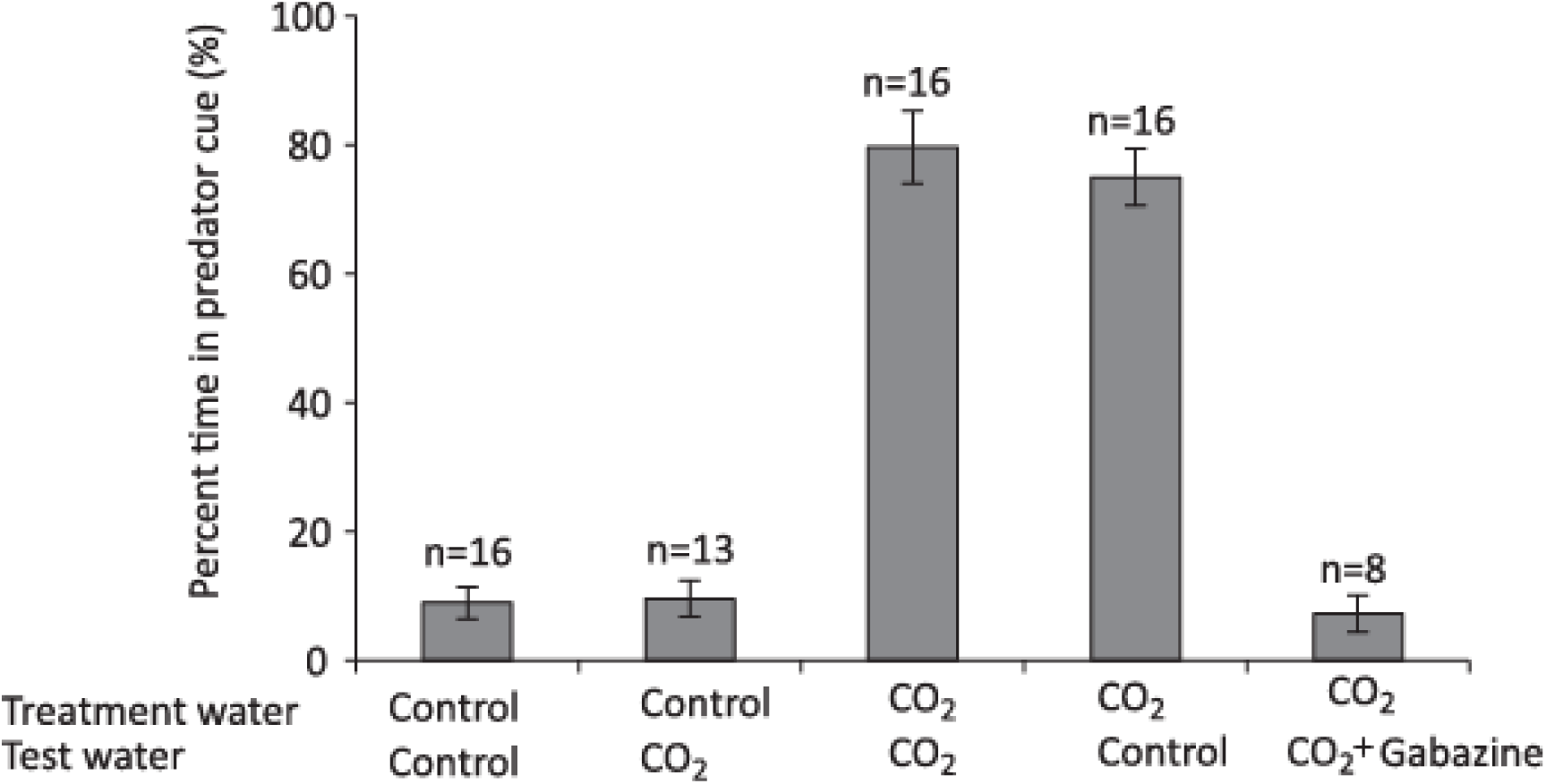
The effect of rearing treatment water (control or elevated CO_2_) and test water used in a two-channel flume (control or elevated CO_2_) on the olfactory response of larval *A. percula* to predator cue. One stream of water in the flume contained seawater with predator cue and the other stream of water had untreated seawater. Shown is the percent time (mean ± s.e.) that fish from each experimental combination spent in the water stream containing predator cue. The final group shows fish that were treated with 4 mg l^−1^ gabazine in high CO_2_ seawater for 30 minutes before testing in the flume. Sample size shown above the bars.

As with the response to predator odour, treatment water had a significant effect on escape responses of larval damselfish, but there was no effect of test water. Latency to respond significantly increased in fish reared in elevated CO_2_, with fish displaying latencies that were 30% slower compared with fish reared in control water (Figure 2a: *F*_1.14_= 5.971, p = 0.018). There was no effect of test water on latency to respond (p = 0.7). Escape distance following stimulation was significantly shorter for fish reared in high CO_2_ compared with control water (Figure 2b: *F*_1,14_= 6.034, p = 0.02) and there was no effect of test water (p = 0.8). Response speed and maximum speed responded similarly with both kinematic variables declining in fish reared in high CO_2_ compared to fish reared in control water (Figure 2c; *F*_1,14_= 8.678, p = 0.01 and Figure 2d; *F*_1,14_ = 9.015, p = 0.009). Again, there was no effect of test water (p = 0.7 and p = 0.5).

**Figure 2.**
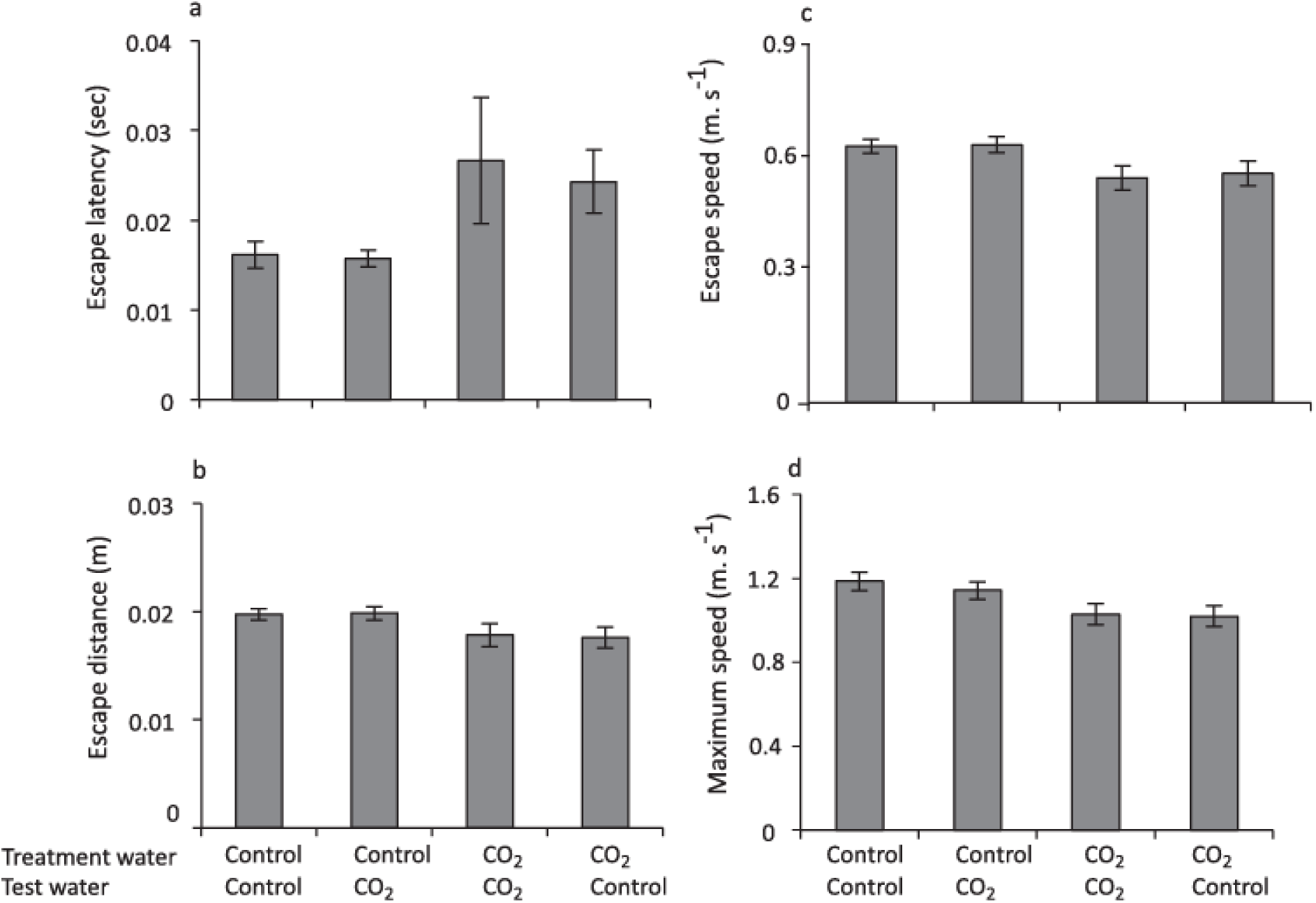
The effect of rearing treatment (control or high CO_2_) and test water used in the experimental trial (control or high CO_2_) on the escape performance of larval *P. amboinensis*. Variables displayed are: (*a*) latency, (*b*) distance, (*c*) mean speed, and (*d*) maximum speed. Errors are standard errors. N = 26, 28, 26, 28 for control - control, control - high CO_2_, high CO_2_ - high CO_2_ and high CO_2_ - control, respectively.

Gabazine reversed the effects of rearing in high CO_2_ (Figure 1). Fish reared in high CO_2_ and then treated with gabazine exhibited a strong avoidance of predator cue, with the response almost identical to fish reared and tested in control water (p=0.96).

## Discussion

Previous studies have demonstrated that CO_2_ levels projected to occur in the surface ocean by the end of this century (700-1000 μatm) impair predator avoidance behaviours in larval reef fishes (reviewed in Nagelkerken & Munday, 2016). However, some of these previous studies have tested the behaviour of high-CO_2_ reared fish using control water in the experimental trials, which could potentially influence the results if there is an immediate physiological response associated with the transfer of fish from high CO_2_ rearing water to control test water. Here we show that the results of these earlier studies are robust and that antipredator behaviour is not affected by using control water in the experimental trials. Larval fish reared in high CO_2_ exhibited the same responses to predator odour in a two channel flume and the same kinematic responses to a startle stimulus regardless of whether high CO_2_ or control water was used in the experimental trial. Similarly, there was no difference in the behavioural response of fish reared in control seawater when tested in either high CO_2_ or control water. These results demonstrate that the choice of test water has not influenced the results of previous laboratory studies into the effects of high CO_2_ on antipredator behaviour.

Our results are consistent with previous pilot studies showing that the test water (control or high CO_2_) used in a two-channel flume did not alter the behavioural response of larval fishes to predator odour (Dixson et al., 2010; Munday et al., 2010). Furthermore, our study extends these findings to the kinematics of predator avoidance, by showing that the type of water used in the test arena does not affect vital components of the escape response, including latency to respond, escape speed and distance travelled. Together these results show that antipredator behaviours of larval fishes that have been reared in high CO_2_ are not compromised by a return to control seawater for experimental testing. These findings are also relevant to field-based studies that have tested the effects of high CO_2_ on reef fishes in their natural habitat (e.g. Munday et al., 2010, 2012; Devine et al., 2012, 2013; Ferrari et al., 2011a; McCormick et al., 2013; Chivers et al., 2014a). Such studies are critical for predicting the impacts of high CO_2_ on reef fish populations; however, they involve rearing larval fish for 4-5 days in high CO_2_ conditions before transferring them to ambient CO_2_ conditions in the field. These studies have demonstrated mortality rates from predation are much higher in juvenile fish that have been exposed to high CO_2_ compared with fish exposed to ambient control conditions (Munday et al., 2010, 2012; Ferrari et al., 2011a; Chivers et al., 2014a). Our results indicate that the results of such studies are unlikely to have been affected by a change in antipredator behaviour due to transplanting fish from high CO_2_ conditions in the laboratory to ambient CO_2_ conditions in their natural habitat.

In contrast to the absence of a test water effect on antipredator behaviours, we found that rearing treatment water had a highly significant effect on antipredator behaviour. Consistent with previous studies (Dixson et al., 2010; Munday et al., 2010), larval clownfish reared in high CO_2_ exhibited a reversal from strongly avoiding predator cue (<10% of time in predator cue in the flume) to a strong attraction to predator cue (nearly 80% of time in predator cue in the flume). Similarly, larval damselfish exposed to high CO_2_ for 4 days exhibited a slower response to a threat stimulus, a slower escape speed and shorted distance travelled compared with fish kept in ambient control water, as has been reported previously (Allan et al., 2013, 2014). Thus, our results provide further confirmatory evidence for the effects of high CO_2_ on antipredator behaviour in larval reef fishes. The differences in escape speed and escape distance between high CO_2_ and control fish were relatively small in the escape response trial. However, when combined with a much longer latency to respond (>50% increase in time to respond) these changes in kinematic responses would likely have a detrimental effect on the probability of a prey fish escaping a predator attack.

Altered behaviour of fish in a high CO_2_ environment appears to be due to the effects of acid-base regulation on the function of GABA-A receptors (Nilsson et al., 2012; Hamilton et al., 2013; Heuer & Grosell, 2014). Fish prevent plasma and tissue acidosis in high CO_2_ by accumulating HCO_3_^−^ in exchange for Cl^−^ (Brauner & Baker 2009). The GABA-A receptor is a gated ion channel that conducts HCO_3_^−^ and Cl^−^ when activated by the neurotransmitter GABA. The altered concentration of HCO_3_^−^ and Cl^−^ in fish exposed to high CO_2_ could decrease or reverse the flow of these ions through the GABA-A receptor, causing a reversal in receptor polarization (Heuer & Grosell, 2014). Evidence for the role of GABA-A receptors in behavioural impairment of fish in high CO_2_ has come from studies showing that treatment with gabazine, a GABA antagonist, reverses the effects of high CO_2_ on behaviour (Nilsson et al., 2012; Chivers et al., 2014a; Chung et al., 2014, Lai et al., 2015; Ou et al., 2015; Regan et al., 2016). In the first of these studies, Nilsson et al. (2012) found that gabazine reversed the impaired response to predator odour exhibited by clownfish that had been reared in high CO_2_. However, in that study gabazine was administered in control seawater and high-CO_2_ reared fish were tested in control water. Therefore, the effects of gabazine in restoring the response of larval fishes to predator odour has not been previously tested in high-CO_2_ conditions. Here we show that treatment with 4 mg l^−1^ gabazine completely reverses the abnormal behavioural of high-CO_2_ fish to predator odour, restoring their natural aversion to predator odour. This contributes to a large number of independent studies pointing to a key role of GABA-A receptors in the behavioural impairment of fish exposed to high CO_2_.

While we found no effects of using control or high CO_2_ water in experimental trials to test the effects of high CO_2_ on fish behaviour, Sundin and Jutfelt (2016) reported differences in relative lateralization of juvenile goldsinny wrasse when tested in control water versus high CO_2_ water after 21 days in high CO_2_. Their study did not include the reverse treatment, where control reared fish were tested in high CO_2_ water, and there is the potential for learning effects in control fish because the same fish were tested on three different occasions, at days 9, 19 and 21. Nevertheless, that study does suggest that the response of some behavioural traits could be sensitive to differences in the test water used in behavioural trials. Further experiments with a broader range of behavioural traits and using experiments that are not complicated by the repeated testing of the same individuals are required to test this possibility.

Pioneering studies into the effects of high CO_2_ on fish behaviour used control seawater in experimental trials due to logistical constraints and concerns about the possible effects of low pH on olfactory cues. Furthermore, pilot studies indicated that there was no difference in the behaviour of fish that had been exposed to high CO_2_ when tested in control or high CO_2_ water. Our results confirm the results of those initial studies by showing that test water (control of high CO_2_) had no effect on the antipredator responses exhibited by fish reared in either control or high CO_2_ conditions. Nevertheless, we do not advocate the continued use of control seawater for testing the behavioural effects of high CO_2_ in marine organisms. Methods for treating seawater to replicate future pCO_2_ levels are now sufficiently validated that it is possible to treat most experimental arenas with the respective treatment water that the animals have been reared in, and there may be some behavioural traits that are sensitive to different test water in the experimental trials. Therefore, most reliable and unambiguous results will likely come from using the respective treatment water in experimental trials testing the effects of high CO_2_ on the behaviour of marine organisms.

## Acknowledgments

We thank staff at JCU’s Marine and Aquaculture Facility and the Australian Museum’s Lizard Island Research Station for logistical support. This research was funded by the Australian Research Council Centre of Excellence for Coral Reef Studies and a Yulgilbar Foundation Fellowship (S.-A.W.).

